# *Pseudomonas aeruginosa* MucP contributes to RNA phage resistance by targeting phage lysis

**DOI:** 10.1101/2024.12.09.627492

**Authors:** Hee-Won Bae, So-Yeon Kim, Shin-Yae Choi, Hyeong-Jun Ki, Se-Jeong Ahn, You-Hee Cho

## Abstract

Lytic phages culminate their lifecycle by causing lysis of the infected host cell. Despite extensive research on the molecular mechanisms of phage lysis, our understanding of anti-phage resistance mechanisms during the lysis stage remains less understood. Here, we demonstrated that MucP, a site 2 protease of *Pseudomonas aeruginosa* (PA), mitigates the activity of the RNA phage PP7 lysis protein (LP), which contains a transmembrane (TM) helix, suggesting that MucP act as a resistance mechanism against PP7-induced lysis. We identified an LP variant (LP*) having enhanced helical propensity due to P26L and S40L mutations, which was unaffected by MucP and exhibited killing activity against PA strains that are resistant to the wild type LP, with an inverse correlation between MucP activity and LP susceptibility. A PP7 mutant with LP* exhibited MucP-escaper phenotype such as discernable plaque formation on MucP-expressing cells. These results suggest that MucP targets the RNA phage LP at the TM helix in certain strains, providing a resistance function compromising phage lysis by utilizing an existing bacterial enzyme in PA.

**IMPORTANCE:** Despite the importance of cell lysis as the last stage of phage lifecycle, the factors influencing coordinated lysis remain elusive in the context of phage-bacteria interactions. Our study identifies MucP, a membrane protease in *Pseudomonas aeruginosa* (PA), as a resistance factor against the RNA phage PP7 by destabilizing its lysis protein (LP), with lower helical tendency at transmembrane (TM) domain. An LP variant with higher helical tendency exhibits strong killing activity against PA isolates, revealing an inverse correlation between MucP activity and LP susceptibility. Since MucP is a conserved protease in mucoid conversion of PA, we propose that MucP offers an intrinsic or passive defense or resistance mechanism against the RNA phage, whose activities vary among the diverse PA isolates.

## INTRODUCTION

The phage lifecycle generally consists of five distinct steps: phage adsorption, genome entry, material synthesis, virion assembly and phage release [1]. As the final stage of the lifecycle, host cell lysis follows the phage virion assembly for all phages except for the filamentous phages that release the progeny phages without cell lysis [2, 3]. As more and more phage particles are assembled inside the cell, the cell reaches a certain point, where it can no longer contain the increasing number of virions and eventually undergoes cell lysis [4]. The phage-mediated cell lysis is one of the most critical events to shape the bacterial population in the ecosystems, which contributes to restructuring bacterial community for fitness management and to releasing cellular matters for nutrient cycling [5, 6].

This coordinated process is crucial for phage fitness during the interaction with the bacterial hosts and thus tightly regulated by the phage lysis systems, which will vary depending on the complexity of the phage species. Two key lysis proteins (holins and endolysins) are typically produced to facilitate cell lysis [2]: holins are membrane proteins that are associated with the bacterial cytoplasmic membrane, forming lesions in the membrane, whereas endolysins are the enzymes that degrade the bacterial peptidoglycan. As a third and new component, spanins are often found in the phages infecting Gram-negative bacteria, since they function in the disruptive linkage of the cell membrane and the outermembrane (OM) [7]. This mode of lysis requiring multiple gene products is exploited in most double-stranded DNA phages. In contrast, some phages of relatively small size (e.g., single-stranded DNA or RNA phages) contain only a single gene for cell lysis, which has been only recently studied compared to the multiple gene lysis systems that have been extensively characterized so far [8, 9]. It is probable that these phages could release the progeny phages only when the bacterial cell membrane is affected by the single gene lysis proteins (LPs) that contain transmembrane (TM) domain and are associated with the cell membrane [10].

The critical contribution of cell lysis to phage fitness implies that this step should be precisely regulated, especially for the phages with a single gene LP. In the present study focusing on the LP of the RNA phage, PP7, which infects some *Pseudomonas aeruginosa* (PA) strains, we revealed that the dysfunction of a site 2 protease, MucP, enhanced the susceptibility to PP7, but not other DNA phages. We observed that the PP7 LP was significantly destabilized by MucP due to the lower- helical propensity residues (P26 and S40) in the TM helix. And an inverse correlation between MucP activity and LP susceptibility was observed in various PA clinical isolates. Furthermore, LP* with higher helical propensity mutations (P26L and S40L) exhibited the MucP-resistance and thus exhibited killing activity against the PA strains with high MucP activity, suggesting that another role of MucP as a phage resistance or as an intrinsic or nonprofessional defense to disturb the RNA phage-mediated host cells lysis in PA strains.

## RESULTS

### LP-mediated cell lysis does not occur in PP7-resistant strains

As an initial attempt to monitor the LP-mediated cell lysis in PA strains, we developed inducible expression systems for RNA phage LP proteins. As shown in Figure. 1A for PP7 LP, we used the *P_tac_*promoter to drive the isopropyl-β-D-thiogalactoside (IPTG)-inducible expression of the LP genes, which were chromosomally integrated at the *att*Tn*7* site [11]. Nucleotide substitutions were introduced at the ribosome binding sites and the start codon to optimize LP expression in PA strains. Additionally, we created fusions of LP with the fluorescent protein, mNeonGreen at either the N-terminus (mNG-LP) or the C-terminus (LP- mNG) and confirmed their comparable killing activity in PAO1 (Figure 1B). mNG-LP was used to monitor PAO1 cell lysis under a fluorescent microscope (Figure 1C and Movie S1). We clearly observed membrane association followed by dispersive concentration of mNG-LP proteins, leading to cell lysis. This observation supports the role of PP7 LP as a membrane holin most likely forming multimers during the host cell lysis.

**Figure 1.**
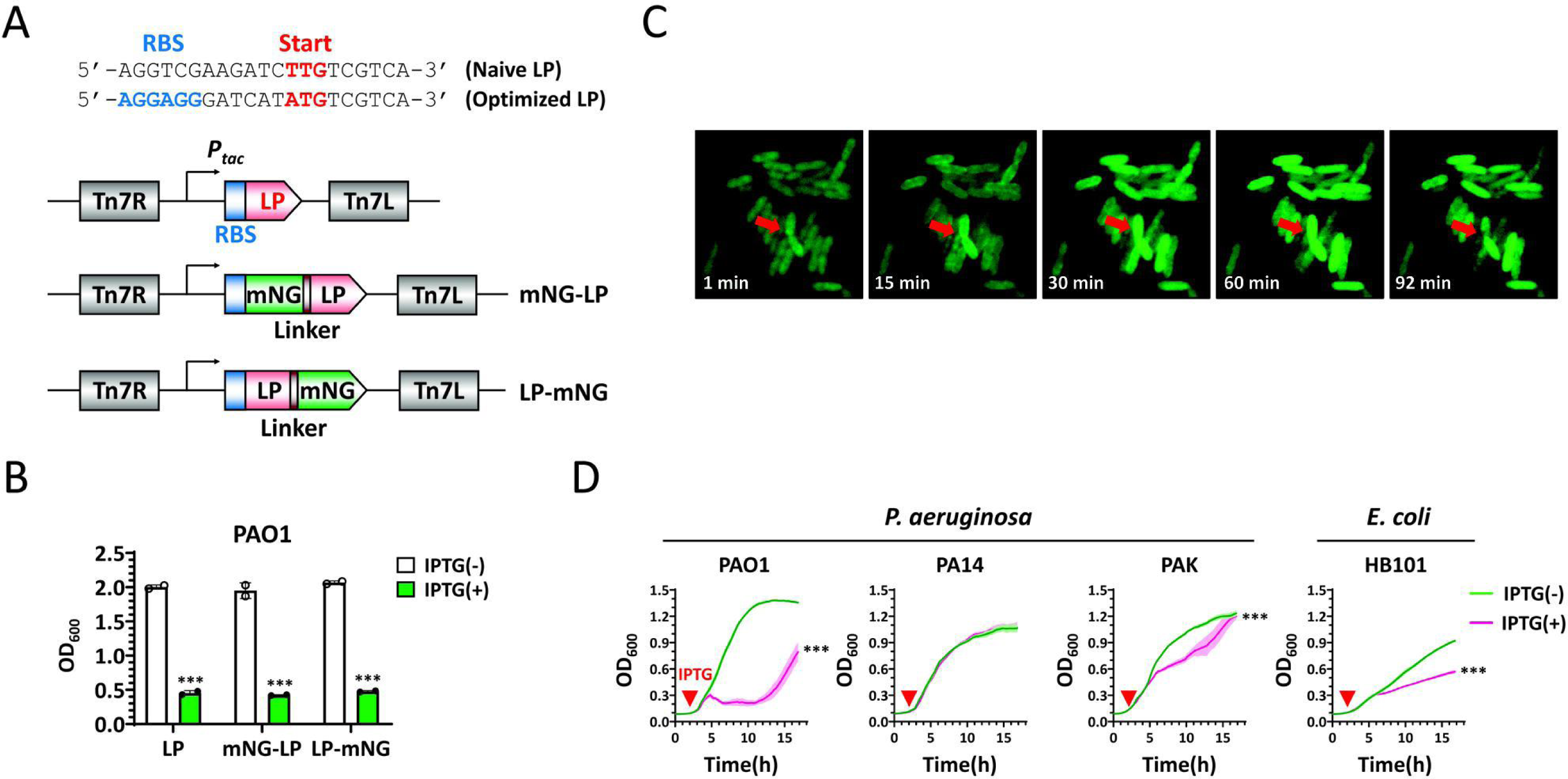
Bacterial killing mediated by PP7 LP expression. A. Schematic representation of LP expression systems. The LP genes with or without linker-fused mNeonGreen (mNG) fluorescent protein at the N- or C- terminal end (mNG-LP or LP-mNG, respectively) were chromosomally integrated at the *att*T7 site under the inducible *P_tac_*promoter. Optimized ribosome binding site (RBS) and the start codons (ATG, bold) are shown alongside the naïve LP gene with the TTG start codon. B. Growth inhibition of PAO1 by LP variants. PAO1 cells expressing LP, mNG-LP, or LP-mNG were grown for 5 h with (+) or without (-) IPTG induction and subjected to the OD_600_ measurement. Statistical significance based on paired *t*-test (one-tailed *p* value): ***, *p* < 0.001. C. Microscopic observation of LP-mediated cell lysis. PAO1 cells with mNG-LP were grown for 3 h, treated with 1 mM IPTG for 10 min, placed on a 1mM IPTG-containing agarose pad, and observed by fluorescent microscopy for 152 min (Movie S1). Representative images are shown from Movie S1 with the corresponding time points. Arrows highlight membrane localization and cell lysis. D. LP-mediated growth inhibition of PA strains and *E. coli* HB101. PAO1, PA14, PAK, and HB101 cells with the LP expression system were grown in 96-well format liquid culture for 18 h with IPTG induction (arrowhead) at 140 min and the growth was monitored by measuring the OD_600_ every 20 min for 17 h. Statistical significance based on paired *t*-test (one-tailed *p* value): ***, *p* < 0.001.

To investigate the role of LP-mediated cell lysis in the PP7 lifecycle, we constructed an LP-free PP7 mutant (Y19) using the cDNA system [12]. As shown in Figure S1A and B, the TAT codon of Y19 was changed to the ochre stop codon (TAA) by inserting A to minimize potential reversion by nucleotide substitution, considering the higher reversion in RNA viruses [13]. This insertion leads to the 2 tandem ochre stop codons (Figure S1B). We found that the Y19 phenotypes differed in plaque morphology and plaque formation efficiency: they formed very tiny and turbid plaques and ∼4 log reduction in plaque formation efficiency even at the same particle amount (Figure S1C) More importantly, the discernable larger plaques were due to the reversion as demonstrated in Figure S1D. These results suggest that LP-mediated cell lysis plays a crucial role in the phage lifecycle and phage fitness with the spontaneous mutations in LP highly restricted.

Given the host spectrum of PP7, which infects some group II pilin-containing PA strains such as PAO1 [14], we investigated whether LP-mediated host cell lysis occurred in other strains. As shown in Figures 1D and S2A, PP7 LP exhibited differential killing activity against three widely used PA strains: no killing activity against PA14, the intermediated killing activity against PAK, and complete killing activity against PAO1. Notably, expression of PRR1 LP resulted in the complete cell lysis in all three strains (Figure S2B). The MS2 LP was active only against *Escherichia coli* strain, HB101 that was susceptible to PP7 LP (Figure S2C). These results indicate that PA14 and PAK may have host barriers limiting PP7 LP activity. Alternatively, PAO1 may possess factors facilitating PP7 LP activity that are absent or not fully functional in PA14 and PAK.

### The *mucP* mutants show higher susceptibility to PP7

Given the two possibilities, we initially performed random transposon mutagenesis of PA14 with the mNG-LP expression system to identify potential RNA phage resistance mechanisms that interfere with the LP-mediated lysis. A total of 33,408 transposon insertion mutants were evaluated for IPTG-induced cell lysis, as described in method details. Three clones (14G5, 15H9, and 21E4) exhibited IPTG-induced cell lysis (Figure S3), corroborating that some factors might interfere with LP-mediated lysis during the phage lifecycle. We mapped the transposon insertion sites using arbitrary PCR, as previously described [15]. We selected 14G5 for further analysis, due to its strong lysis phenotype with the transposon insertion mapped at the *mucP* gene, a site 2 protease involved in mucoid conversion of PA strains (Figure 2A and B). It should be noted that MucP is a core gene in PA, suggesting that the phenetic differences involving MucP could account for the variation in LP-mediated lysis between PAO1 and PA14. To verify this, we created the in-frame *mucP* deletions in both PAO1 and PA14P (a PP7-suceptible PA14 derivative; 15) to assess if their susceptibility to PP7 differed from the parental strains. As shown in Figure 2C, PP7 formed more discernible and larger plaques on the *mucP* mutant than on the wild type in PA14P, compared to PAO1. Notably larger plaques also formed on the *mucP* mutant in PAO1. In contrast, the plaque size was significantly reduced by ectopic expression from the multi-copy plasmid containing the *mucP* gene in both PAO1 and PA14P. These MucP effects were also confirmed in liquid culture (Figure 2D) and were not observed for DNA phages (MP29 and MPK7), suggesting that the MucP-mediated effect is specific to the PP7 LP.

**Figure 2.**
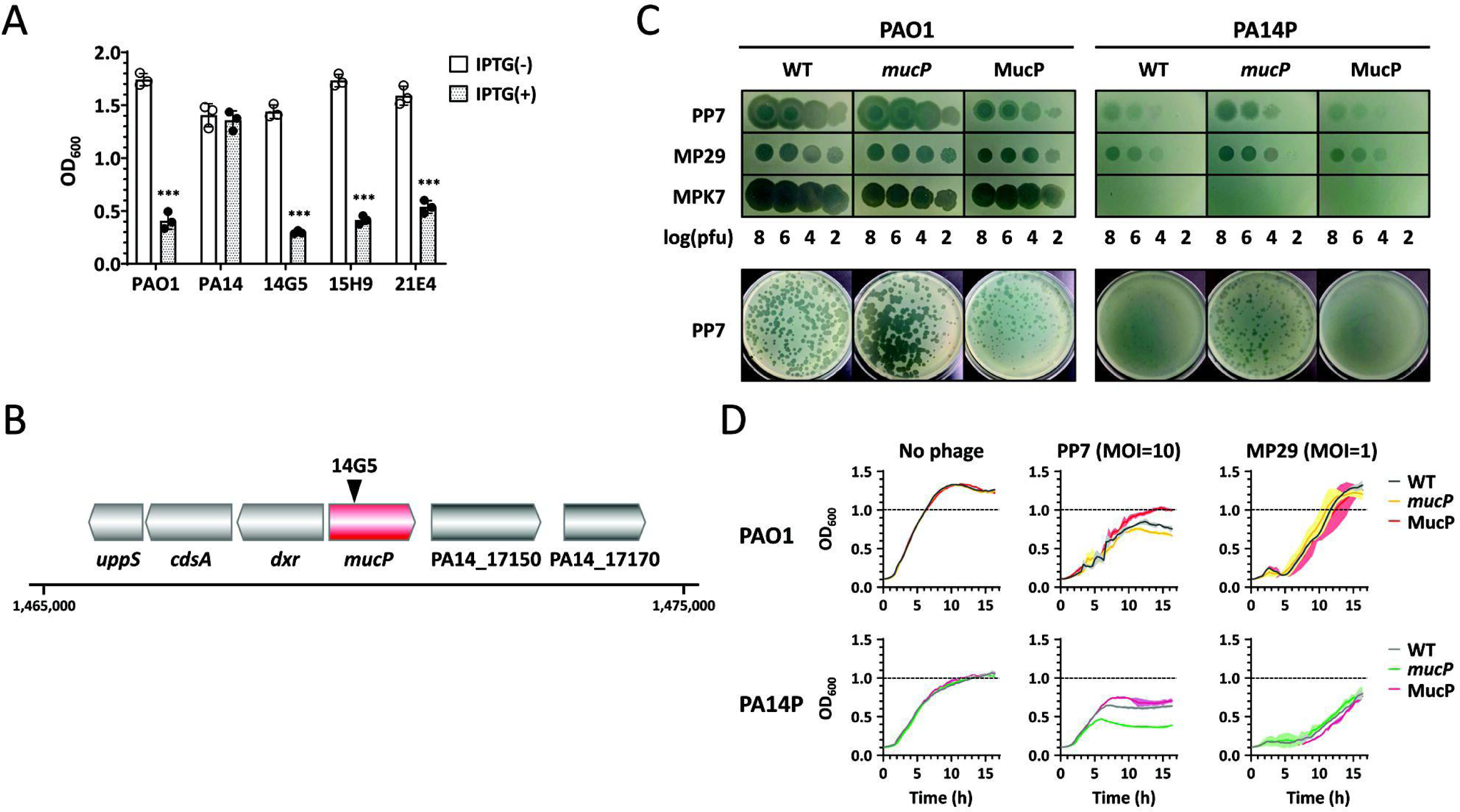
Identification of the genes for LP resistance. A. Growth inhibition of the transposon mutants. Transposon mutant (14G5, 21E4, and 15H9) cells were grown in 15-ml tube format liquid culture for 3 h with (+) or without (-) IPTG induction. PAO1(LP) was included as the control (PAO1) and PA14(LP) is the parental strain (PA14). The growth was assessed by OD_600_ measurement. Statistical significance based on paired *t*-test (one-tailed *p* value): ***, *p* < 0.001. B. Mapping of transposon insertion in 14G5. Schematic representation of the genomic region containing the transposon insertion region is shown with the PA14 genome coordinate indicated. The transposon insertion site is designated by an arrowhead in the *mucP* gene. C. Spotting and plaque phenotypes of the *mucP* deletion mutants. Phage spots from four serially 100-fold diluted phage samples of PP7, MPK7, or MP29 and their plaques were investigated on the bacterial lawns of the wild type (WT), the isogenic *mucP* mutants (*mucP*) and the WT with the plasmid-borne *mucP* gene (MucP) of PAO1 (left) and PA14P (right). D. Growth curve of the bacterial cells in 96-well format liquid culture. Culture suspensions of the WT, *mucP*, or MucP of PAO1 (upper) and PA14P (lower) were mixed with the phage lysates of PP7 (at MOI of 10), MP29 (at MOI of 1) in LB medium and then incubated for 15 h. The growth was assessed by measuring the OD_600_ at every 20 min for 16 h.

### MucP destabilizes PP7 LP

The observation that the PRR1 LP induces cell lysis in PA14 led us to hypothesize that there might be specific determinants differentiating the PRR1 and PP7 LPs in relation to the MucP function. Given that both LPs possess transmembrane (TM) helices and that MucP is a site 2 membrane protease, it is likely that the TM helices of these LPs exhibit different characteristics. Upon examining the 23 amino acid sequences in the TM domains of both LPs and the MS2 LP, we found that the PP7 LP contains lower helical propensity amino acids compared to the PRR1 and MS2 LPs (Figure 3A).

**Figure 3.**
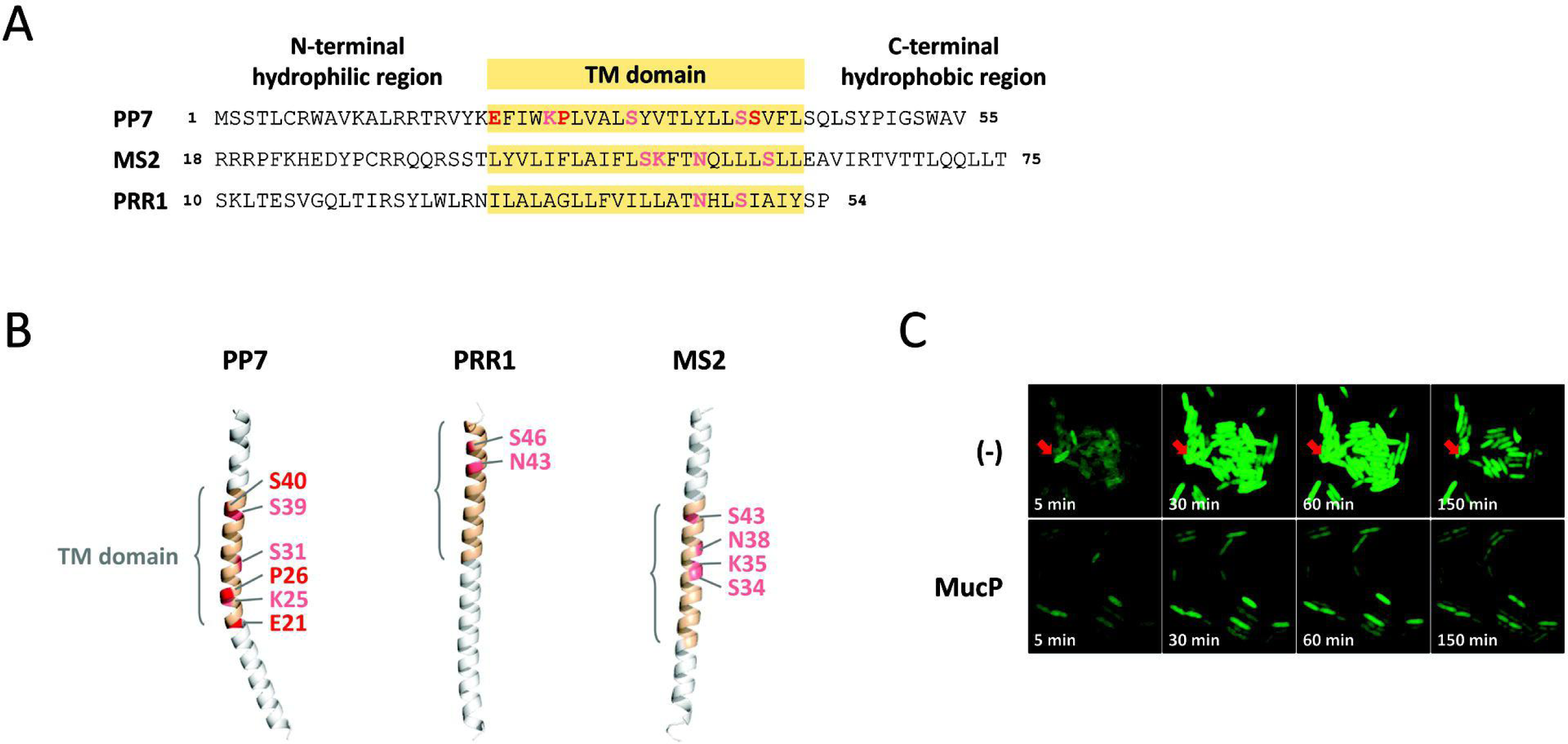
Sequence and predicted structure of the LP transmembrane domains. A. Multiple alignment of LP proteins. Schematic representation of the 3 regions of LP proteins is designated: N-terminal hydrophilic region, transmembrane (TM) domain, and C-terminal hydrophobic region. The 55-amino acids of PP7 LP are shown in alignment with the partial sequences of PRR1 and MS2 LPs at the TM domain (in yellow shade). The low helical-propensity amino acids in a membrane environment are designated in bold. B. Structural prediction of the LP proteins from PP7, PRR1, and MS2. Structures were predicted using AlphaFold2. Transmembrane domains and the low helical- propensity amino acids are highlighted. C. Microscopic observation of the LP stability by MucP overexpression. PAO1 cells with mNG-LP with either pUCP18 (-) or pUCP-mucP (MucP) grown for 3 h was treated with 1 mM IPTG for 10 min, transferred to a 1 mM IPTG-containing agarose pad, and then observed under fluorescent microscope. Representative images up to 150-min observation are shown for each with the corresponding time points indicated. The arrows indicate the cell lysis in the absence (-) of plasmid-borne MucP expression.

Low helical propensities in the membrane or non-polar environments are typical for charged or polar amino acids, with proline having the lowest helical propensity due to the structural rigidity imparted by its unique cyclic side chain structure [16]. Figure 3A and B shows that the PP7 LP are richer in low helical propensity amino acids than the other LPs. AlphaFold2 modeling of each LP monomer also revealed a slightly kinked structure only in the PP7 LP around the proline (P26) (Figure 3B). It is also remarkable that low helical propensity amino acids are necessary for efficient cleavage by site 2 membrane proteases [17, 18]. These findings suggest that the PP7 LP would be a better substrate for site 2 membrane proteases, such as MucP, compared to the other LPs. To verify this, we used the mNG-LP expression system in PAO1 to determine if MucP could destabilize the PP7 LP. As shown in Figure 3C, fluorescence from mNG-LP almost disappeared, accompanied by a lack of cell lysis upon ectopic expression of MucP.

### Helical propensity of the LP TM domain influences MucP-susceptibility

To further verify the relationship between TM helical propensity and MucP susceptibility, we introduced single and multiple point mutations to increase hydrophobicity, minimizing amino acid changes in the overlapping RNA replicase (RP) gene (Figure S4A). As shown in Figure S4B, four LP mutants (E21V, P26L, S31I, and S40L) demonstrated higher killing activity against PAO1 than the wild type LP. Notably, only P26L exhibited killing activity against PA14, indicating that P26 is indeed crucial for LP instability in PA14. We then created two double mutants (E21VP26L and P26LS40L) and one triple mutant (E21VP26LS40L) to test their killing activity against PA14 (Figure S4B and 4A). The triple and the P26LS40L mutants showed enhanced killing activity against PA14, and importantly, their activity was not compromised by the ectopic expression of MucP in PA strains. This suggests that both P26 and S40 contribute to LP helical instability, with P26 being the major contributor. Structural modeling of the P26LS40L protein (designated as LP* hereafter) showed a straighter TM helix (Figure 4B).

**Figure 4.**
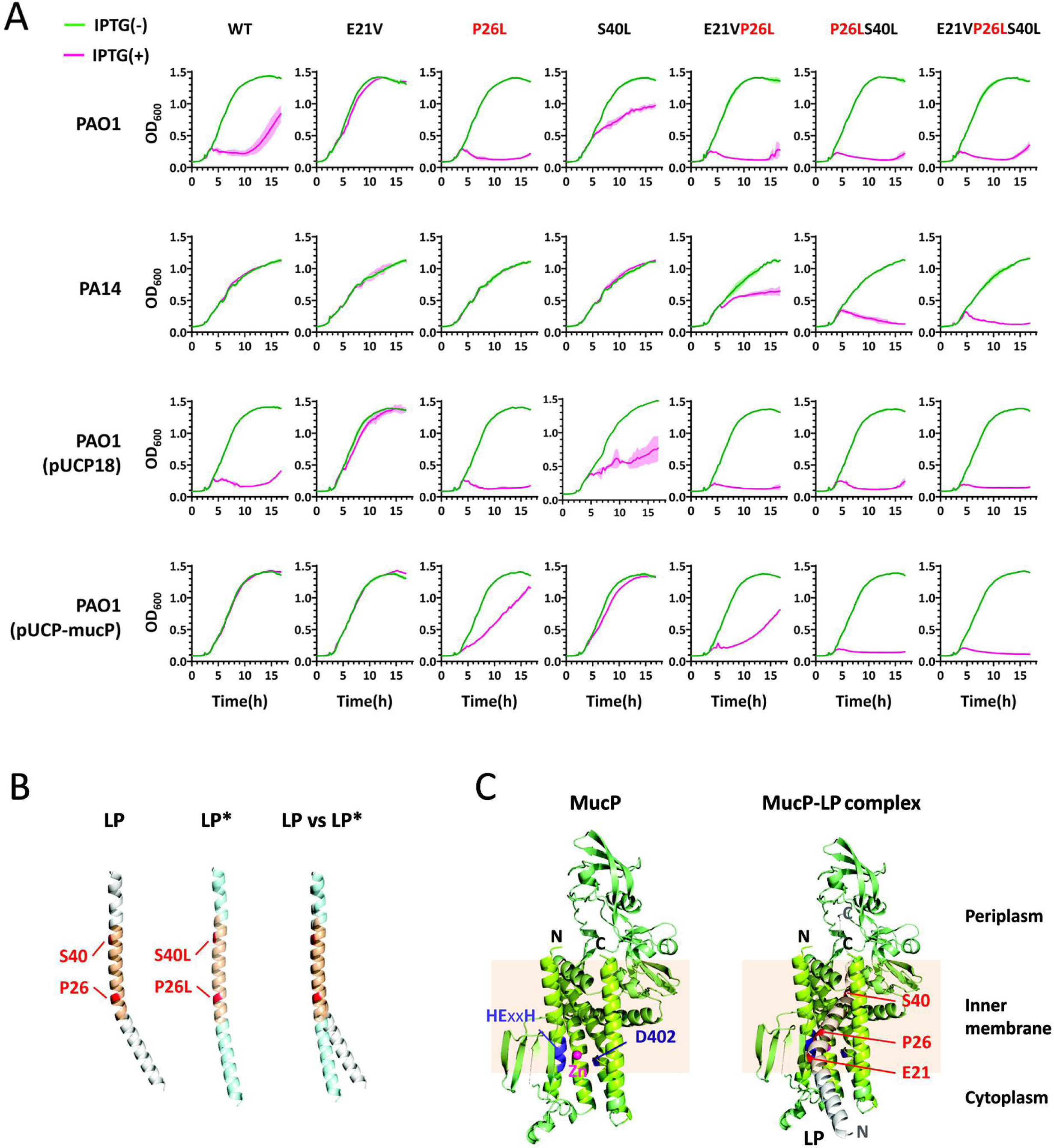
Bacterial killing mediated by expression of TM helix mutants. A. Growth inhibition of PA strains. PA (PAO1 and its derivatives or PA14) cells with the expression system for each TM helix mutant (as designated at the top) were grown in 96-well format liquid culture for 18 h with IPTG induction at 140 min and the growth was monitored by measuring the OD_600_ every 20 min for 17h. PAO1 derivatives are PAO1 overexpressing MucP (PAO1(MucP)) and its vector control (PAO1(pUCP18)). B. Structural prediction and superimposition of the LP (gray) and LP* (skyblue). Structures were predicted using AlphaFold2. TM domains as well as the two key residues low helical-propensity amino (P26 and S40) for LP vs high helical- propensity amino acids (L26 and L40) for LP* acids are designated. C. Predicted structural modeling of MucP and the MucP-LP interaction. The predicted interaction interface involves the close association between the catalytic site of MucP (HExxH) and the P26 at the TM domains in the inner membrane. The N-terminal hydrophilic region of LP is protruding at the cytoplasm, whereas the short C-terminal region of LP is at the periplasm.

The AlphaFold2 model of the PP7 LP and MucP complex revealed that the MucP catalytic site, characterized by the HExxH zinc metalloprotease motif, and P26 are closely located at their contact interface, where Zn^2+^ can be coordinated presumably by H21, E22, H25, and D402 (Figure 4C) [19]. The helical axis appears significantly distorted at the end of the TM domain in this model, supporting that P26-induced lower helical propensity of PP7 LP contributes mainly to MucP susceptibility and differential killing activities in PAO1 and PA14. Largely based on this modeling, we suggest that the PP7 LP could be the direct substrate of MucP, although the functional multimeric states of the PP7 LP monomers and the subsequent conformation of the membrane-disrupting structures of the LP multimers need to be more comprehensively elucidated.

### Clinical strains with higher MucP activity exhibit higher LP resistance

To explore if LP* enhances the killing spectrum against diverse PA strains, we conducted a growth inhibition assay using 28 PA strains as shown in Figure 1D confirmed that all but one strain (PMM50) was killed by LP* upon IPTG induction, demonstrating the enhanced activity and/or broadened killing spectrum of LP* (Figure S5).

We then correlated the differential LP susceptibility with MucP activity by measuring the basal *algD* transcript levels via RT-qPCR, as increased MucP activity elevates the *algD* transcription during mucoid conversion of the PA strains [20]. Figure 5A showed the basal *algD* expression levels during the aerobic liquid culture clearly differed in the 28 strains, with higher transcription in PA14 and PAK than that in PAO1. A strong inverse correlation (*r*^2^=0.692) was found between *algD* transcription (i.e., MucP activity) and LP-mediated growth inhibition at ∼13 h in the 28 strains (Figure 5B), indicating that higher MucP activity results in greater LP resistance. Only one outlier (PMM50) existed, presumably due to other contributing factors, in that the LP-mediated lysis might involve complicated mechanisms yet to be unveiled that occur in the cytoplasmic membrane. Nevertheless, the significant correlation between LP susceptibility and MucP activity in clinical PA strains suggests that MucP-mediated LP resistance is an intrinsic resistance mechanism against PP7 and related RNA phages in PA strains.

**Figure 5.**
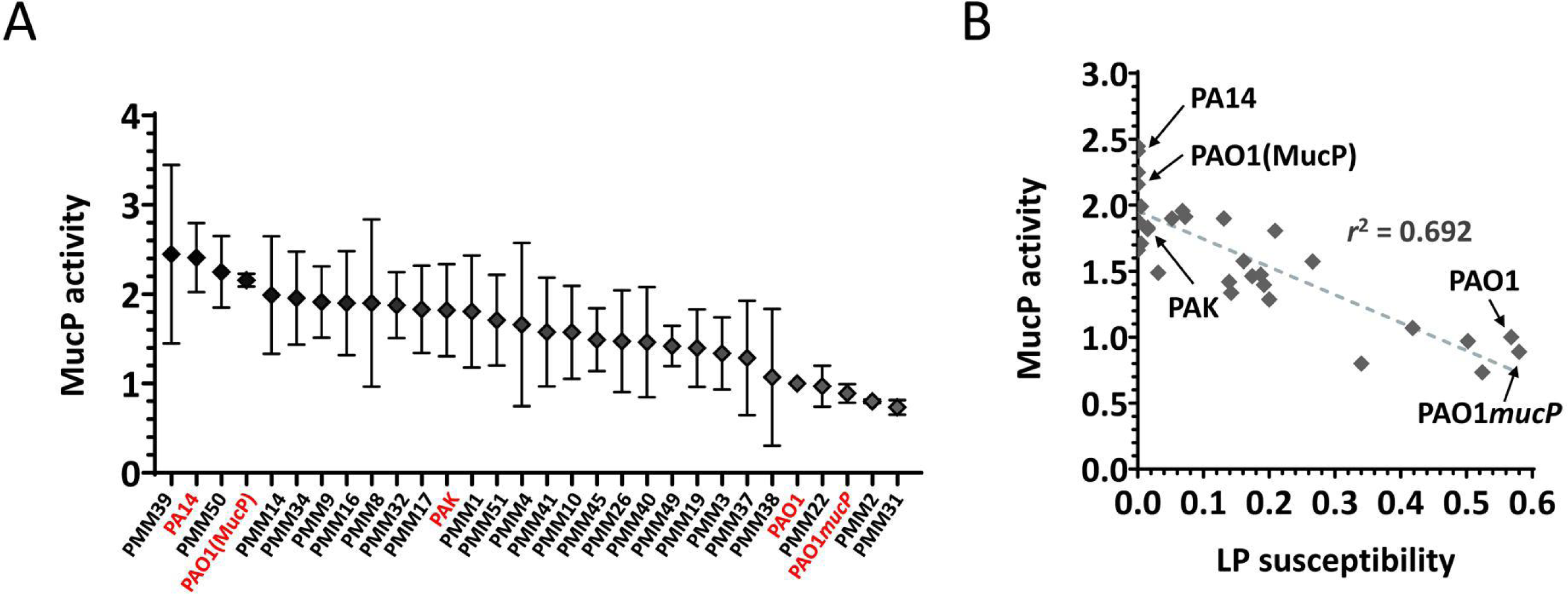
MucP activity and LP susceptibility in clinical strains. A. MucP activity in clinical strains. MucP activity as assessed by the *algD* transcript level relative to the *rpoA* one in the clinical strains indicated. PAO1 and its derivatives, PAO1 *mucP* and PAO1(pUCP-mucP), are included for comparison and indicated. PA14 and PAK are also indicated. B. Correlation between MucP activity and LP susceptibility in clinical strains and PAO1 derivatives. MucP activity values from panel **A** are plotted over the corresponding LP susceptibility values as calculated from Figure S5 (see Methods). PAO1 and its derivatives, PAO1 *mucP* and PAO1(pUCP-mucP) as well as PA14 and PAK are included for comparison and indicated (in red).

### PP7 variant with LP* can form clear plaques even on MucP-expressing cells

Discovery of LP* prompted us to question how LP* could affect the lifecycle of PP7 by changing the lysis physiology in phenotypically diverse PA strains involving differential MucP activity. To assess the involvement of LP* activity in PP7 propagation, we created a PP7 variant (PP7*) possessing LP* by using the cDNA-based reverse genetic system. Infectious PP7* phages were obtained, in that the P26L and S40L mutations in LP* were designed for silent mutations at T3 and L17 of RP, as they occurred at the third codon positions (Figure S4A). As shown in Figure 6A, PP7* could form clearer and more discernable plaques with greater plaque formation efficiency on the MucP-overexpressing PAO1 cells, suggesting that PP7* could be an escaper from the MucP-mediated LP destabilization in PA strains. We also investigated the growth inhibition of PAO1 cells by PP7* in liquid culture (Figure 6B). PP7* displayed greater growth inhibition in the MucP-overexpressing PAO1 cells than the wild type PP7, notably at lower multiplicity of infection (MOI). The wild type PP7 displayed slightly greater growth inhibition in the wild type PAO1 and its *mucP* mutant, clearly at higher MOI (MOI of 10). This result suggests that LP* could resist the MucP-mediated resistance in PA strains with higher MucP activity, with the PP7* phage showing the escaper phenotype from MucP-mediated LP destabilization during this RNA phage lifecycle.

**Figure 6.**
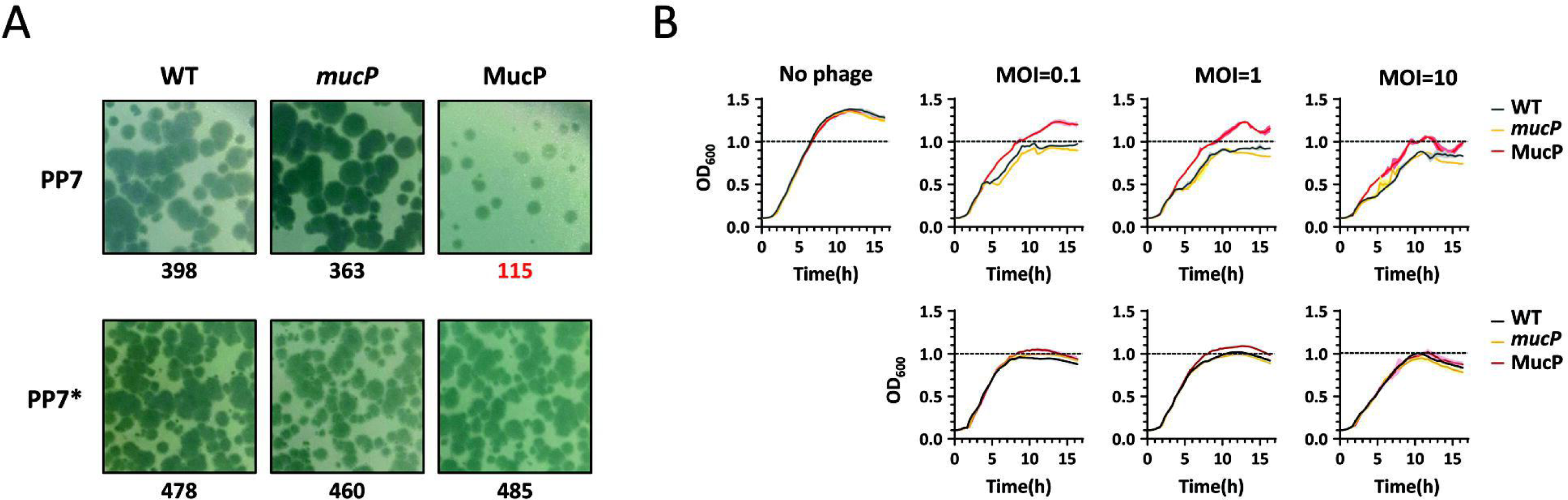
Phenotypes of PP7 phages with LP vs LP*. A. Plaque formation of PP7 phages with LP*. Appropriately diluted phage samples of either PP7 or its derivative with LP* (PP7*) were allowed to form plaques on the lawn cells of PAO1 (WT), its *mucP* mutant (*mucP*) and PAO1 harboring pUCP18-mucP (MucP). The numbers at each bottom indicate the numbers of plaques observed in the entire plates. B. Growth inhibition by PP7*. PAO1 (WT), its *mucP* mutant (*mucP*) and PAO1 harboring pUCP18-mucP (MucP) cells were mixed with either PP7 or PP7* at the indicated multiplicities of infection (MOIs) (0.1, 1, and 10) and then grown in 96-well format liquid culture for 18 h. The growth was monitored by measuring the OD_600_ every 20 min for 17 h. The horizontal lines at OD_600_ of 1.0 are shown for comparison.

## DISCUSSION

One of the fundamental questions in phage biology is what determines the host range or host spectrum of a phage. This is governed by the successful progression through the phage lifecycle stages, overcoming bacterial resistance or defense mechanisms exerted at each of the five distinct lifecycle stages. Our research focuses on the molecular and submolecular determinants of the host range for the small RNA phage (fiersphage), PP7, which uses the group II pilin of the type IV pilus (TFP) as its receptor [14, 21]. This study identifies another bacterial determinant in the host range of PP7: MucP, a membrane zinc metalloprotease known for its role in mucoid conversion in PA by regulating alginate production through the proteolytic degradation of MucA, a membrane-embedded anti-sigma factor [22]. MucP appears to interfere with the LP-mediated cell lysis stage of the phage lifecycle, likely through direct interaction with the PP7 LP. Our previous efforts to identify PA factors affecting the host range of PP7 have shown that the other lifecycle stages can be disrupted by host factors involving the pilin (*pilA*) gene for phage adsorption and a secondary lipid A acylase (*htrB2*) gene for genome entry [14, 15]. These findings suggest that these factors can restrict the phage lifecycle, leading to a failure in proper progeny production of PP7. Consequently, genes like *pilA*, *htrB2*, and *mucP* can be considered as “general” or “intrinsic” phage resistance or defense systems, since they are conserved across all PA strains with other physiological roles not specialized for the defense against phages, given that these genes are part of the PA core genes. Functional alterations, either by allelic variation (as with *pilA*) or by activity alteration (as with *htrB2* and *mucP*) in some strains might direct general resistance against PP7. These resistance mechanisms, which exploit the existing bacterial systems, can be distinguished from the “professional” or “acquired” defense mechanisms that are typically obtained through prophage integration or gene mobilization [23, 24]. For example, a number of known professional defense genes are clustered at the two core defense hot spots in PA genomes, with signatures of mobile genetic elements [25].

This study also highlights that cell lysis, the final stage of the phage lifecycle, can be targeted by bacterial resistance mechanisms, given that host lysis is a temporally scheduled process, with the lysis timing having impact on the phage fitness [26, 27]. In this regard, cell lysis timing should be precisely regulated, which has been verified mostly from DNA phages with multiple gene lysis systems. The concerted involvement of each component from the systems is crucial for the lysis timing control. In general, holins are the lysis timer that mediates cell membrane dysfunction, which is followed by secretion and/or activation of endolysins and by outermembrane disruption by spanins [4, 28]. In contrast, the small DNA or RNA phages with only one LP could have different mechanisms of lysis timing control, with its oligomerization leading to the pore structures that resemble the holins [29, 30]. Given that the small RNA phages could pass through the cell envelope structures from the inside of the cells, the function of LP might be associated with the lysis timing control as for the DNA phage holins.

It is also experimentally verified in this study that the RNA phage LP is indeed crucial to the fitness of the RNA phages, at least in terms of phage propagation. This needs to be more carefully addressed, because only PP7 has the MucP-susceptible LP, unlike other two RNA phages, PRR1 and MS2, tested in this study. Nevertheless, this is in good agreement with the previous study by Adler et al. [31] that one of the multicopy suppressors for the PP7 LP-mediated cell lysis in *E. coli* is RseP. More importantly, RseP, which is a MucP homolog cleaving the RseA anti-sigma factor [32], is active against the PP7 LP not against the MS2 LP. It should also be noted that the RNA phages could be generated without cell lysis, given that RNA phages can be produced from the cDNA-containing surrogate bacteria not normally infected by the generated phages (e.g., PAK for PP7). Nevertheless, the lysis timing control would be beneficial for the ecological symbiosis between phages and bacteria. In this regard, the phage resistance directed by MucP that targets the PP7 LP might interfere with the proper propagation of PP7 in PA strains with higher MucP activity. Higher MucP activity is one of the phenotypic variations in PA clinical strains with different degrees of mucoid conversion-related adaptation in the context of chronic infection. Thus, further studies on the roles of MucP in the RNA phage lysis control are warranted to provide an insight into the molecular interactions that occur at the various stages of phage lifecycle to shape the evolutionary strategies for the arms race between phages and bacteria.

## MATERIALS AND METHODS

### Bacterial strains and culture conditions

*Pseudomonas aeruginosa* (PAO1, PA14, PAK, and PMM clinical strains) and *Escherichia coli* (HB101, DH5α, and SM10) strains and their derivatives were grown at 37°C using Luria-Bertani (LB) broth or onto 2% agar-bacteriological LB (Acumedia) plates. Cetrimide agar (Difco) plates were used to select *P. aeruginosa* (PA) strains. If necessary, antibiotics were amended at the following concentrations (μg/ml): gentamicin (50), and carbenicillin (200) for PA; gentamicin (25) and ampicillin (50) for *E. coli*. Induction by Isopropyl-β-D-thiogalactoside (IPTG) was performed at 150 μM for 96-well plate liquid culture and 1 mM for 15-ml tube format liquid culture.

### Construction of LP expression systems

The LP genes from PP7, PRR1 and MS2 were amplified with the primer pairs (Table 2) using Phusion™ high-fidelity DNA polymerase (Thermo Fisher Scientific). The amplified products were gel-purified and digested with BamHI and PstI (New England Biolabs) and ligated into mini-Tn*7*T-LAC vector with engineered ribosome binding site optimized for translation in PA. To visualize the LP expression, mNeonGreen was fused to LP with a 15-aa linker (GGGGSGGGGSGGGGS) at either the N-terminus (mNG-LP) or the C-terminus (LP-mNG). The LP mutants were generated by SOEing PCR using the primer pairs listed in Table 2.

**Table 1.**
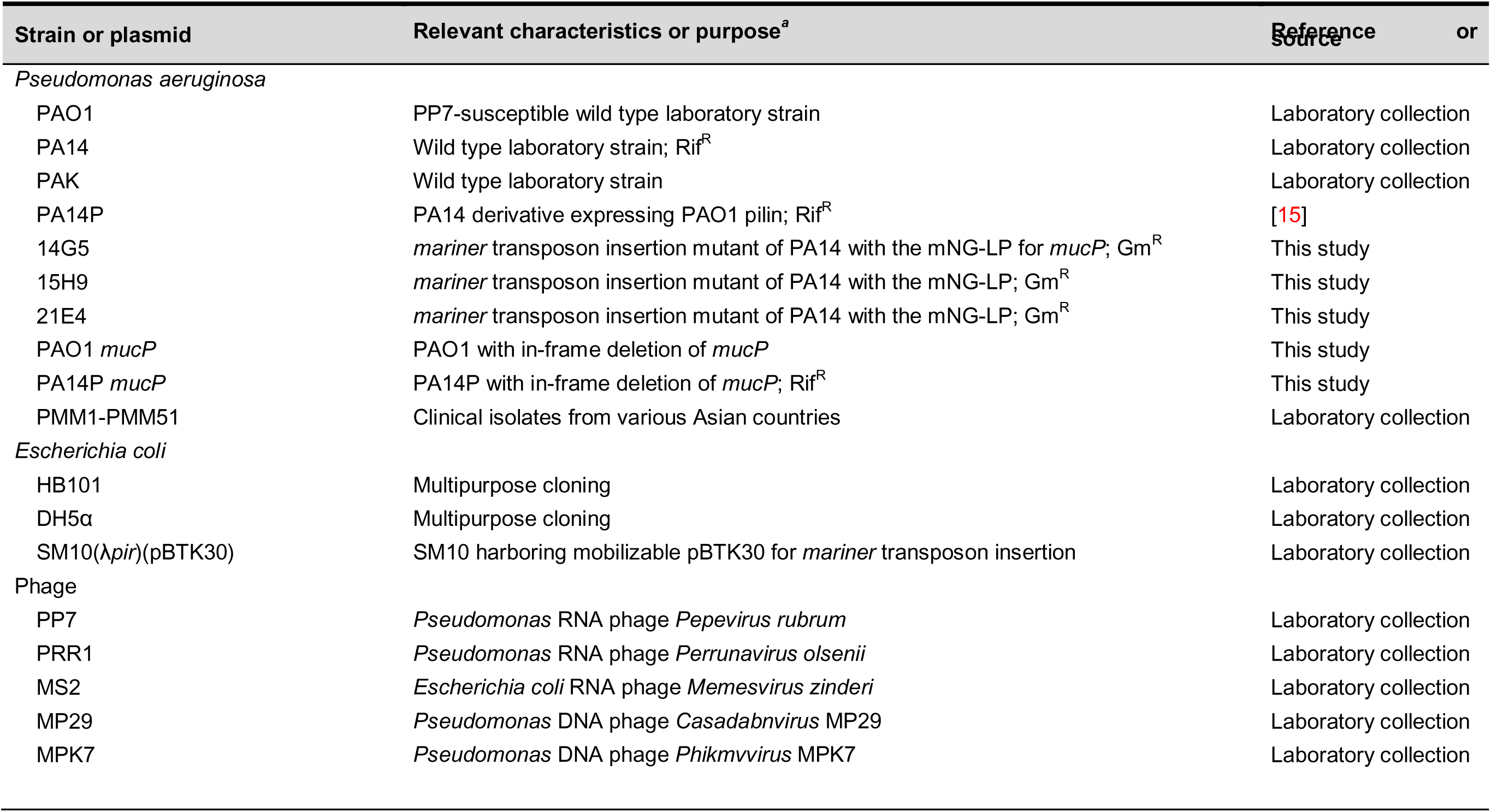
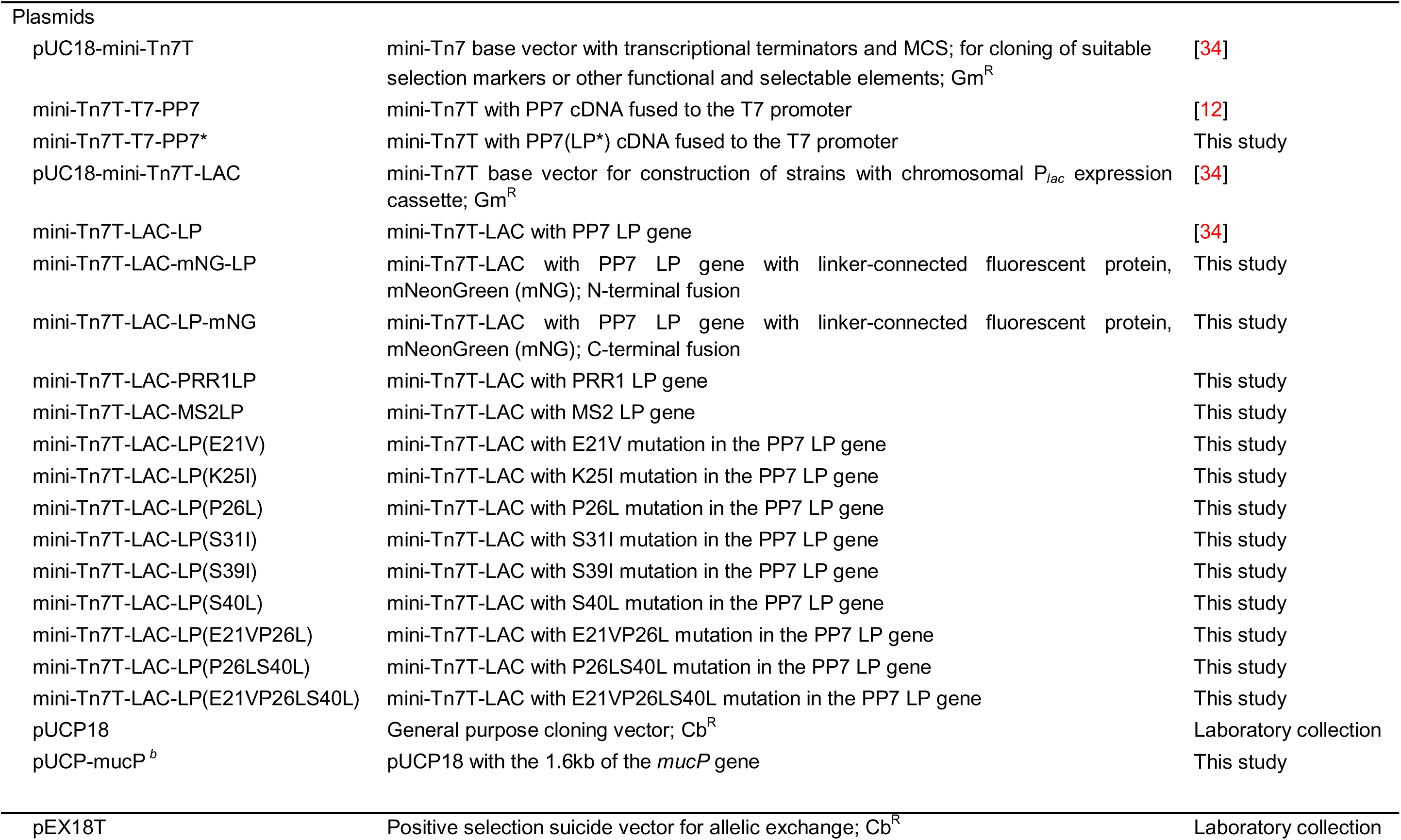
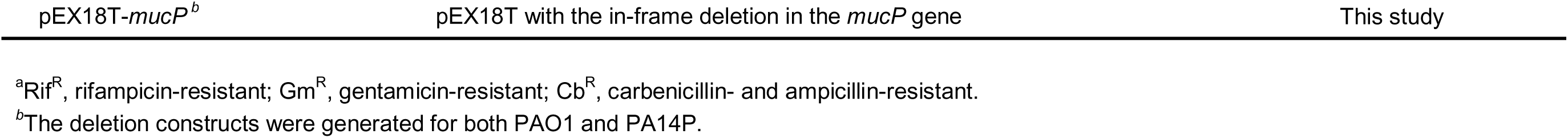
Bacterial and phage strains and plasmids used in this study.

**Table 2.**
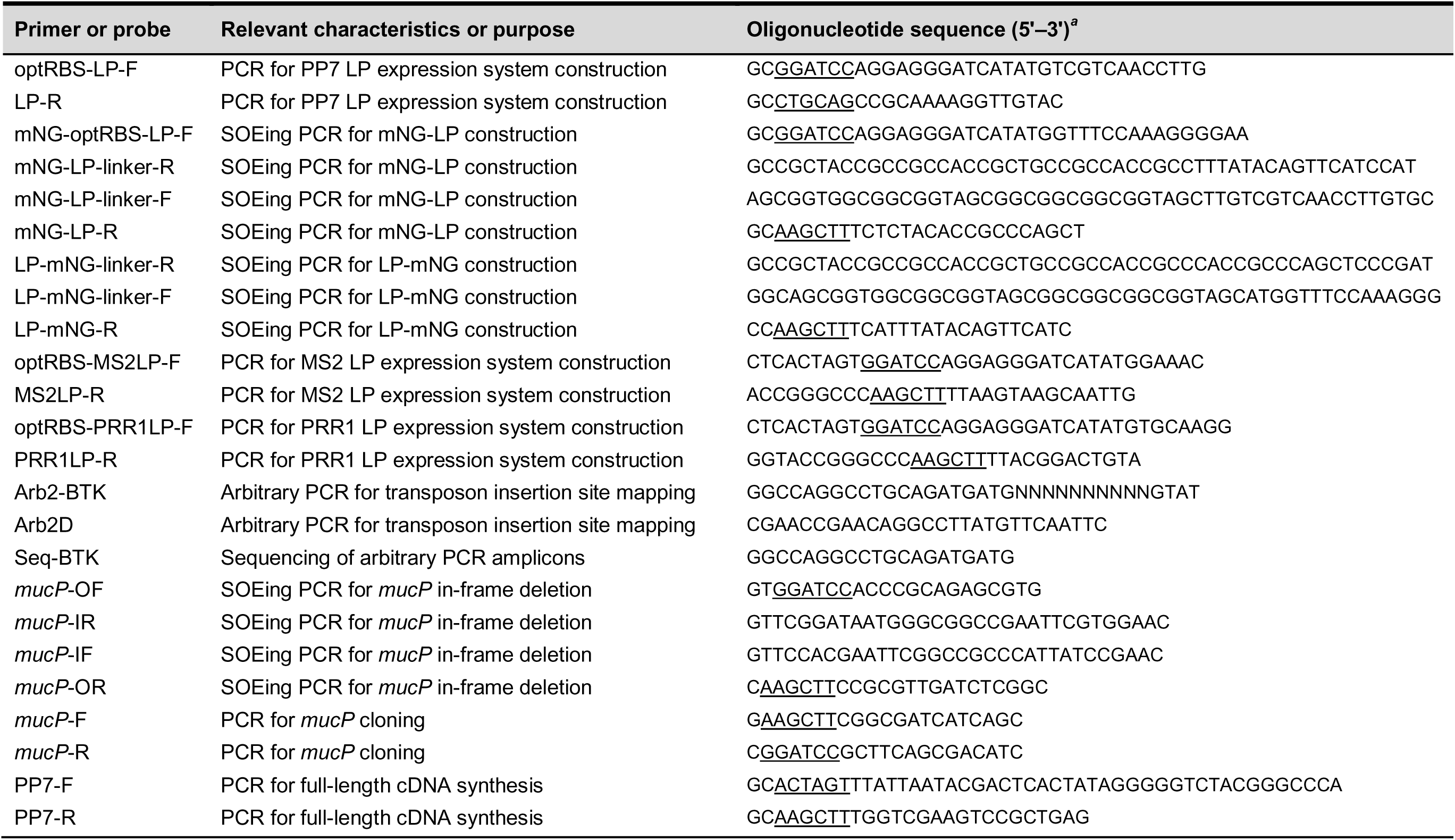
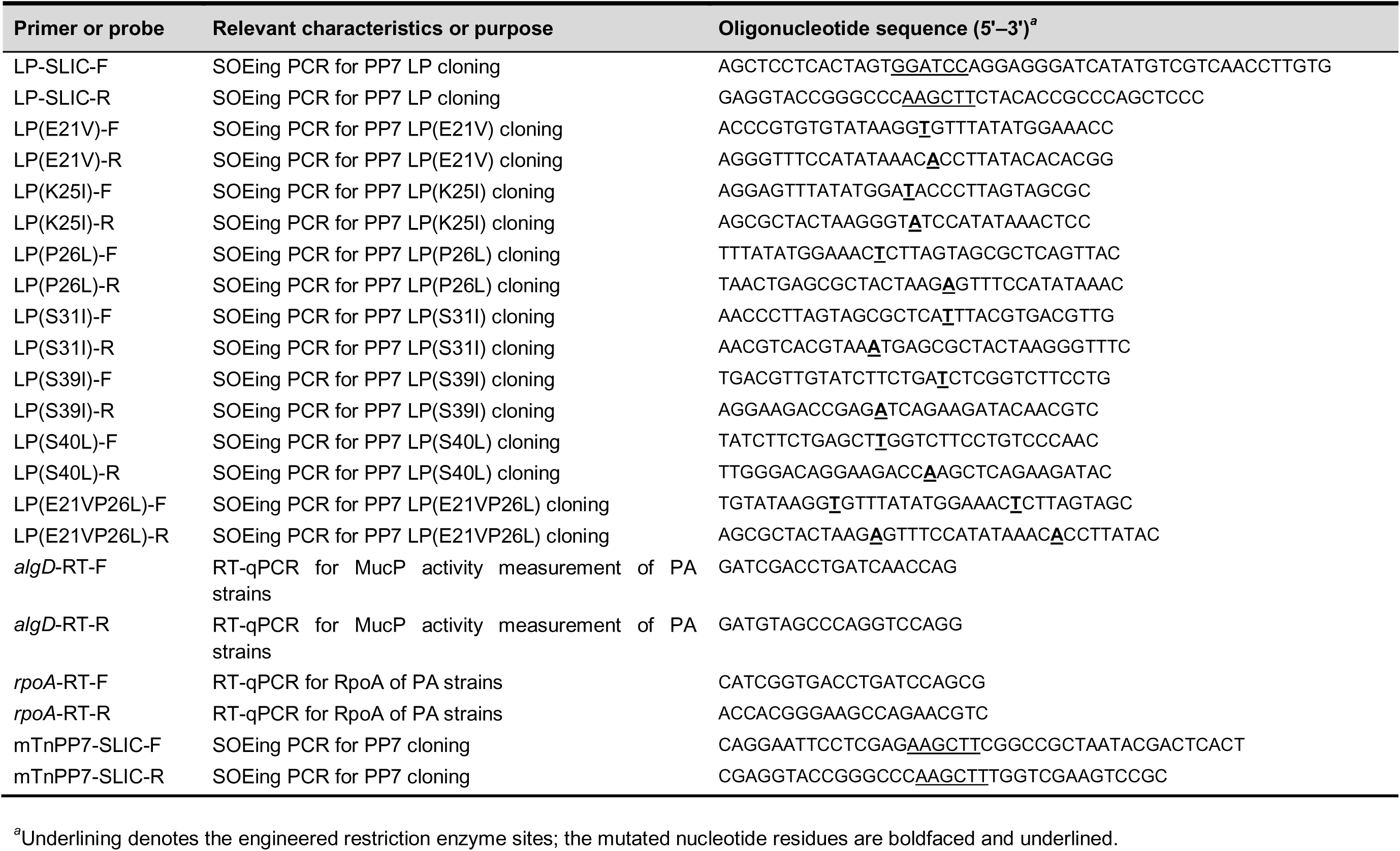
Primers and probes used in this study.

### Fluorescence microscopy

Fluorescence microscopy was done to monitor LP-mediated cell lysis in PAO1. PAO1 carrying mNG-LP was grown to an optical density at 600 nm (OD_600_) of ∼0.2 at 37°C and then treated with 1 mM IPTG. After 10 min, the culture aliquots (1 μl) were transferred onto an agarose pad (2% agarose in M9-citrate minimal medium [0.05% NaCl, 0.1% NH_4_Cl, 1.2% Na_2_HPO_4_, 0.3% KH_2_PO_4_, 2 mM MgSO_4_, 0.1 mM CaCl_2_, and 0.4% citrate] with 1 mM IPTG) and observed under a fluorescence microscope (Zeiss Axio Observer Z1) at 37°C in a temperature controller (LCI, South Korea) up to 3 h. All the fluorescence images were processed by using Zeiss ZEN software.

### LP-mediated killing assay

Bacterial cells were grown overnight in LB broth supplemented with appropriate antibiotics and diluted 1:100 into fresh LB medium. For 96-well plate cultures, 100 μl of each prepared cell was transferred into LB broth in a 96-well plate. The plate was cultured at 37°C with shaking using a microplate reader (BioTek Epoch2) and OD_600_ was measured every 20 min. After 140 min, 50 μl of fresh LB media with or without IPTG (at the final concentration of 150 μM) were added into the culture. OD_600_ measurements were carried out during further incubation for additional 14 h in triplicate.

For test tube cultures, the overnight culture cells were diluted 1:100 into 4 ml of fresh LB medium and grown at 37°C with shaking until they reach OD_600_ of 0.3. Cultures were divided into two groups for treatment by either water or 1 mM IPTG. Cells were grown further at 37°C in a shaker, and the OD_600_ values were measured at the indicated time points.

### Preparation of phage lysates

Phage lysates of MPK7, MP29 and PRR1 were prepared by plate lysate method, using PAO1, while PP7 and MS2 lysates were prepared using PAK cells containing its cDNA that had been cultured in 200 ml LB broth for 16 h at 37°C as described elsewhere [12]. Phage particles were precipitated with 1M NaCl and 10% polyethylene glycol (average molecular weight, 8,000 Da) (Sigma-Aldrich) at 4°C overnight, pelleted by centrifugation, and dissolved in phage buffer (50 mM Tris-HCl [pH 7.5], 10 mM MgSO_4_, 100 mM NaCl). Phage particles were further concentrated by ultracentrifugation at 180,000 × *g* for 6 h and then resuspended in 1 ml of phage buffer. The plaque forming unit (PFU) and the copy number of the phage lysates were determined prior to being used.

### Creation of mutant (Y19 and PP7*) phages

A reverse genetic system for PP7 was used to generate PP7 mutants as previously described [33]. The cDNA for both mutant phages were generated by SOEing PCR (Table 2) and resulting PCR products were cloned into pUC18-mini-Tn*7*T-LAC by SLIC (sequence ligation independent cloning) [34]. The resulting plasmids were introduced into PAK strain by conjugation to generate mutant phage particles.

### Verification and quantification of PP7 phages

Phages were isolated from the plaques and RNA was extracted using TRIzol (Invitrogen). cDNA was synthesized from 10 ng of RNA using SuperScript III reverse transcriptase (Enzynomics) with PP7-R primer. Then, the full-length PP7 cDNA was amplified by PCR using PP7-F and PP7-R primer pair (Table 2). The PCR amplicons were subjected to nucleotide sequencing by a local vender (Macrogen). Measurement of phage copy or virion numbers were performed as described previously [33].

### Phage infection assay

Phage infection was performed either in the liquid cultures or on the plate cultures of PA cells as previously described [35]. For liquid cultures, bacterial growth was assessed spectrophotometrically in the presence of phages at the designated multiplicity of infection (MOI) by measuring OD_600_ using a microplate reader (BioTek Epoch2). For plate cultures, droplets (3 μl) of serially diluted phage samples were spotted onto the lawns of PA cells (100 μl) from the exponential-phase cultures. The plates were incubated at 37°C for 16 h. Alternatively, the phage samples (3 μl) were mixed with 3 ml top agar containing 10^8^ PA cells and then the mixture was plated onto LB agar. Phage plaques were examined after 16 h incubation at 37°C.

### Isolation of transposon mutants

Random transposon mutagenesis was performed by using pBTK30, containing a *mariner*-based transposon [36]. Biparental mating from *E. coli* SM10(λ*pir*) harboring pBTK30 to PA14 with the PP7 cDNA was performed onto LB agar plates for 6 h at 37°C. Selection and counter-selection was done using cetrimide agar plates containing 50 μg/ml gentamicin. A total of 33,408 insertion clones were screened for the mutants exhibiting IPTG-induced cell lysis to isolate 158 primary candidates. After secondary screening, the transposon insertion sites of 3 clones were determined by arbitrary PCR followed by sequencing as described elsewhere [37] using the primers (Table 2).

### Generation of *mucP* and MucP cells

The *mucP*-deleted mutants were created by 4-primer SOEing (splicing by overlap extension) PCR in both PAO1 and PA14P backgrounds (Table 2). The double-crossover mutants were generated by using pEX18T-based allelic exchange as described elsewhere [38] and the deletion mutants were verified by PCR. Also, to confirm the effects by ectopic expression from the multi-copy plasmid containing the *mucP* gene in both PAO1 and PA14P, the *mucP* genes were amplified from PAO1 and PA14 strains using the primers, *mucP*-F and *mucP*-R (Table 2) and the PCR products were cloned into pUCP18 at the BamHI and HindIII sites. The resulting plasmids were introduced into the wild type cells of PAO1 and PA14 by electroporation (Bio-Rad MicroPulser^TM^) [39].

### Prediction of protein structures

The structures of phage and bacterial proteins were predicted using AlphaFold2 (v. 2.3.1). Three models were generated for each, and the best model (ranked_0.pdb) based on the average pLDDT score was selected. The structures were visualized using PyMOL (v. 2.3.0).

### MucP activity assay

Clinical PA strains as well as PAO1, PA14 and PAK cells were inoculated in LB, grown at 37°C to late-exponential phase (OD_600_ of 1), centrifuged at 10,000 × *g* for 2 min. The cells were harvested for RNA extraction using a RNeasy mini kit (Qiagen), according to the manufacturer’s instructions. The RNA samples were subjected to DNase I digestion (Qiagen) and subjected to RT-qPCR using *algD* and *rpoA* primers (Table 2) and the StepOnePlus qPCR system (Applied Biosystems).

### Bioinformatics and statistics

The results were presented as the means with the real data and were analyzed by using GraphPad Prism Version 8.4.3 statistical software. For box and whisker plotting, paired *t*-tests were performed to analyze one-tailed *p*-value with confidence intervals of 95% as described elsewhere [35]. A *p*-value of <0.05 was considered statistically significant.

## ACKNOWLEDGMENTS

This work was supported by the National Research Foundation of Korea (NRF) Grants (NRF-2022R1A2C3003943 and NRF-2022M3A9F3082329)

## AUTHOR CONTRIBUTIONS

H.-W.B. and Y.-H.C. conceived and designed the research. H.-W.B., S.-Y.K., S.-Y.C., H.-J.K., and S.-J.A. designed and performed the experiments, and collected and analyzed the experimental data. H.-W.B. and Y.-H.C. wrote the manuscript. All authors reviewed the manuscript.

---

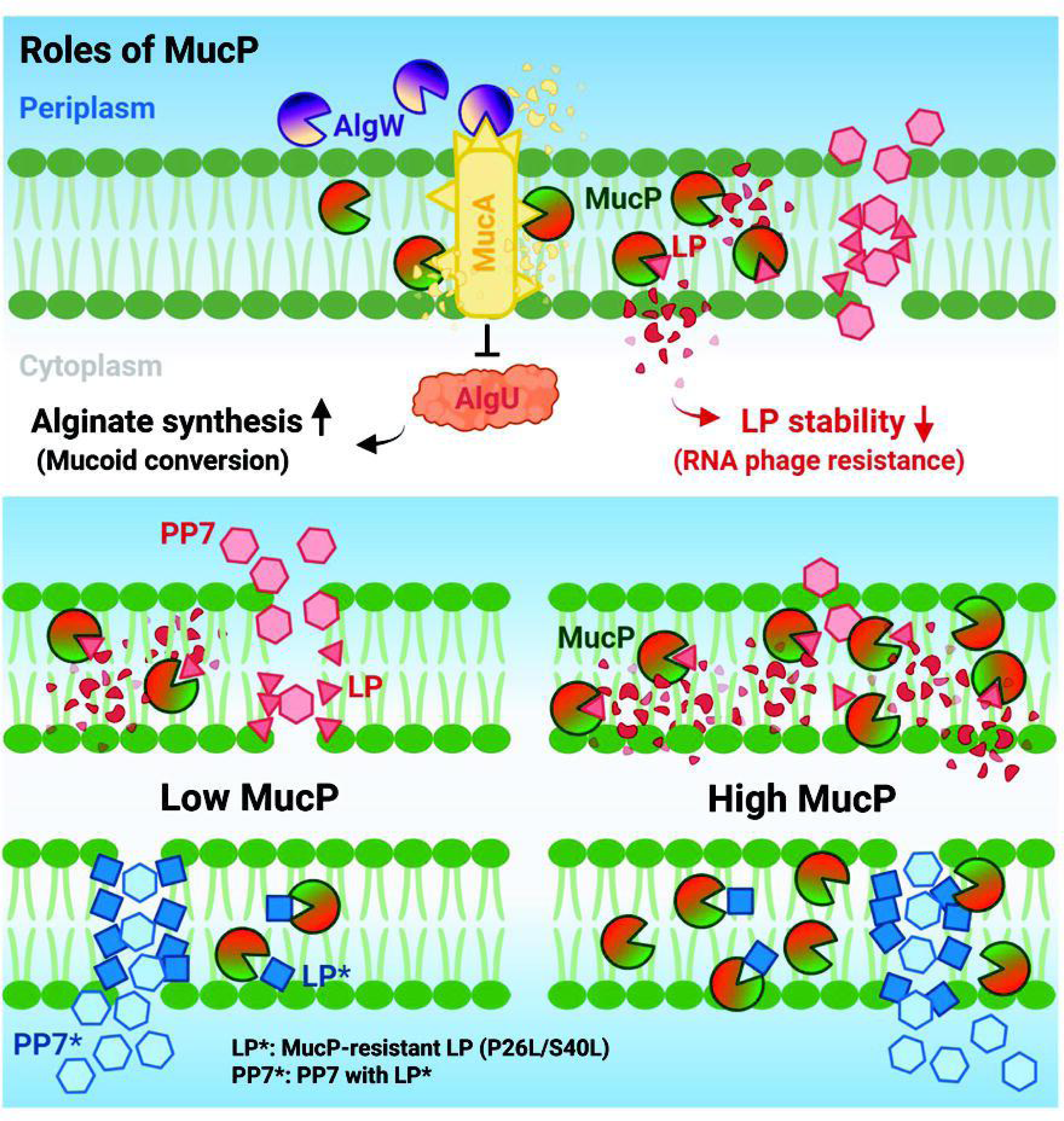

