## Supplementary material for "*Pseudomonas aeruginosa* MucP contributes to RNA phage resistance by targeting phage lysis": SI_Figs

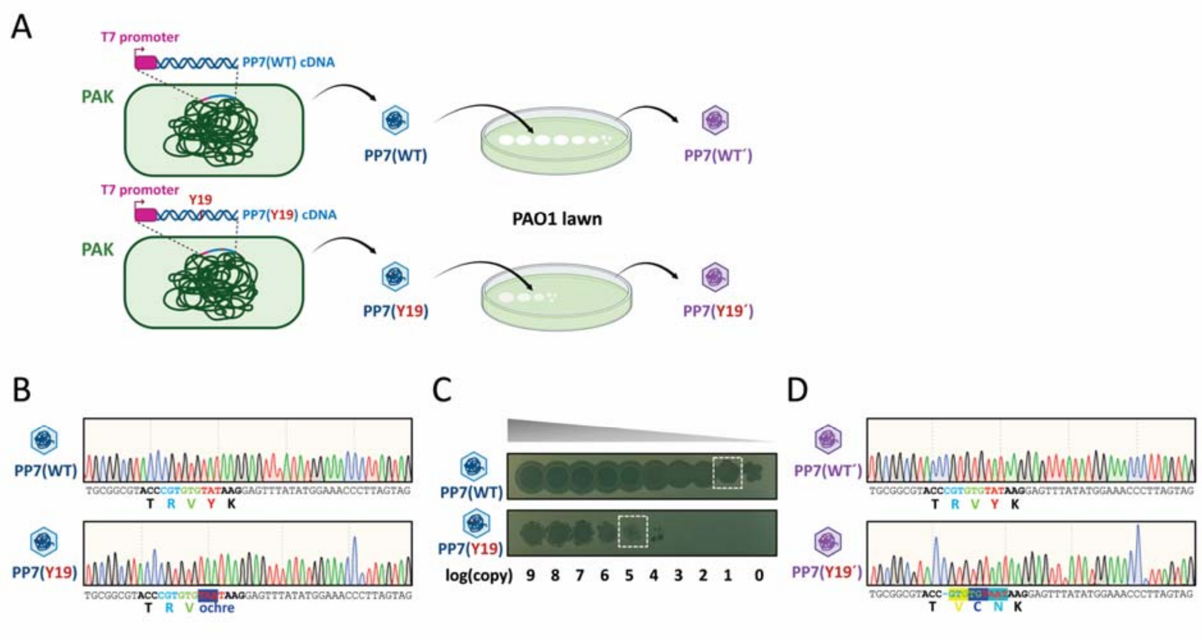


**Figure S1. LP-free PP7 mutant phenotypes,** related to Figure 1

A. Schematic overview of phenotype assessment for the LP-free PP7 mutant (Y19). The cDNA-derived phage particles [PP7(WT) and PP7(Y19)] (Kim et al., 2021) were applied to PAO1 cells to produce progeny phage particles denoted by prime (´) symbols.

B. Sequence verification of cDNA-derived phage particles [PP7(WT) and PP7(Y19)]. Phage RNAs from PP7(WT) and PP7(Y19) were subjected to RT-PCR. The electropherogram displays partial nucleotide sequences of the PCR products, highlighting codons and corresponding amino acids near the TAT codon (in red). In the PP7(Y19) mutant, an “A” insertion changes the 19th tyrosine (Y) codon, TAT to TAAT creating an ochre stop codon (shaded in blue).

C. Plaque formation of cDNA-derived phage particles [PP7(WT) and PP7(Y19)]. Phage samples of PP7(WT) and PP7(Y19) were spotted on PAO1 cells as depicted in A. The numbers indicate the log values of the RNA phage copy numbers. Boxes indicate the plaques selected for sequence verification in **D**.

D. Sequence verification of progeny phage particles [PP7(WT´) and PP7(Y19´)]. Phage RNAs from PP7(WT´) and PP7(Y19´) were subjected to RT-PCR. The electropherogram displays partial nucleotide sequences of the PCR products, highlighting codons and corresponding amino acids as in **B**. in Y19´, a “C” deletion (indicated by “-”) caused a frameshift that bypassed the ochre stop codon, resulting in restoration of LP with three amino acid changes.

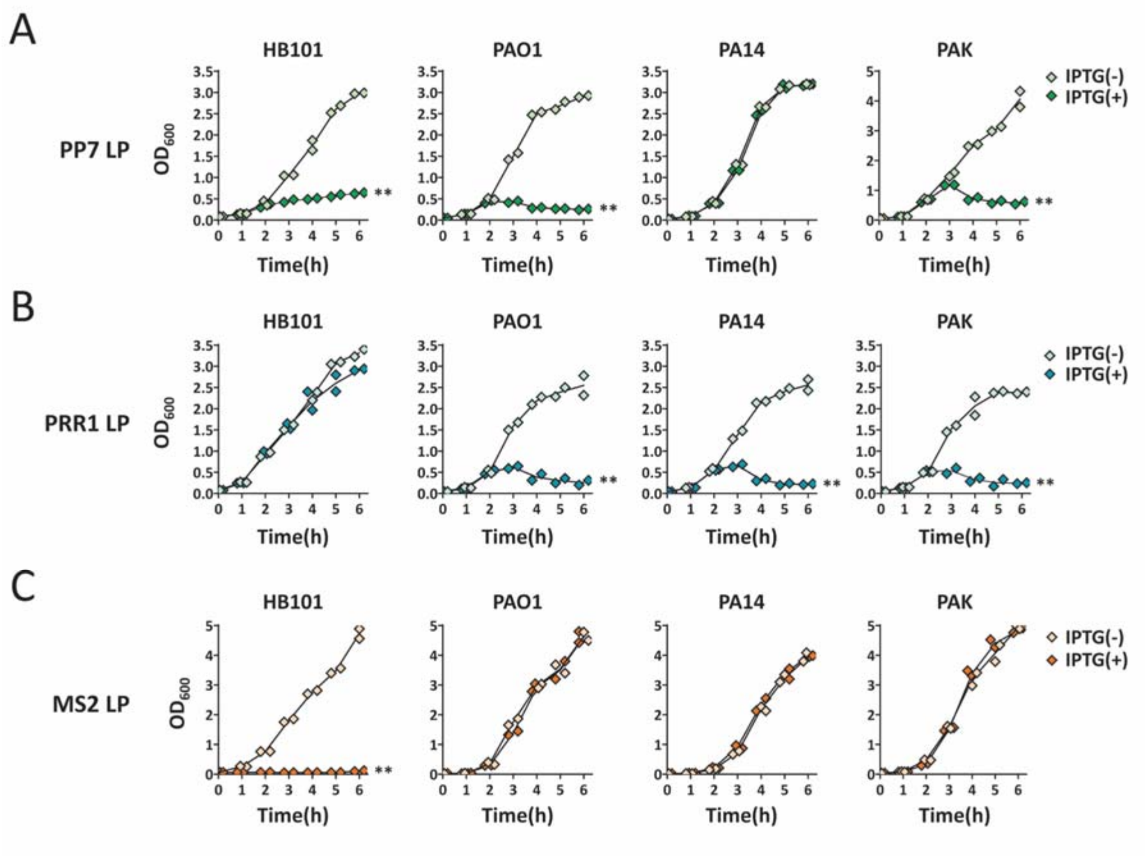


**Figure S2. Bacterial killing mediated by expression of PRR1 and MS2 LP,** related to Figure 1

A-C. PAO1, PA14, PAK, and HB101 cells with the LP expression systems for PP7 (A), PRR1 (B) or MS2 (C) were grown in 15-ml tube format liquid culture for 6 h with (+) or without (-) IPTG induction and the growth was monitored by measuring the OD600 every 1 h for 6 h. Statistical significance based on paired t-test (one-tailed p value): ns, not significant; **, $p < 0.005$.

**Movie S1. Microscopy of LP-mediated cell lysis,** related to Figure 1

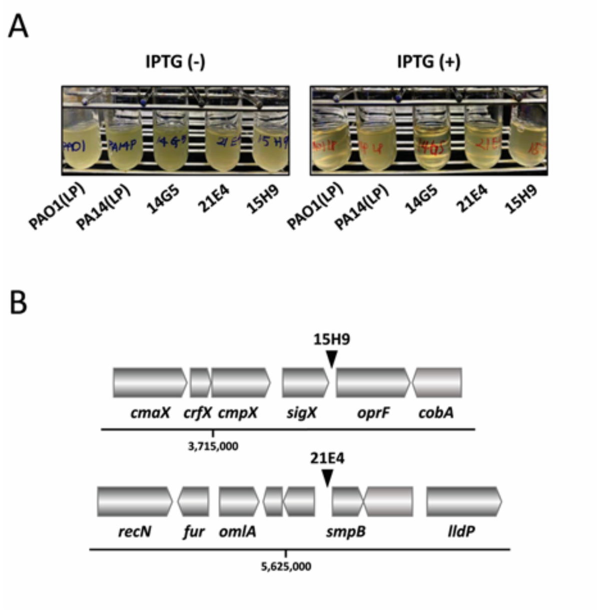


**Figure S3. Identification of PA14 transposon mutants with LP-susceptibility,** related to Figures 1 and 2

A. Growth inhibition of the transposon mutants. Transposon mutant (14G5, 21E4, and 15H9) cells were grown in 15-ml tube format liquid culture for 6 h with (+) or without (-) IPTG induction. PAO1(LP) served as control and PA14(LP) as the parental strain. The growth inhibition was monitored by visual inspection.

B. Mapping of transposon insertions in 15H9 and 21E4. Schematic representation of the genomic regions containing the transposon insertion sites is shown with the PA14 genome coordinate indicated. The insertion sites for 15H9 and 21E4 are designated by arrowheads in the intergenic regions designated.

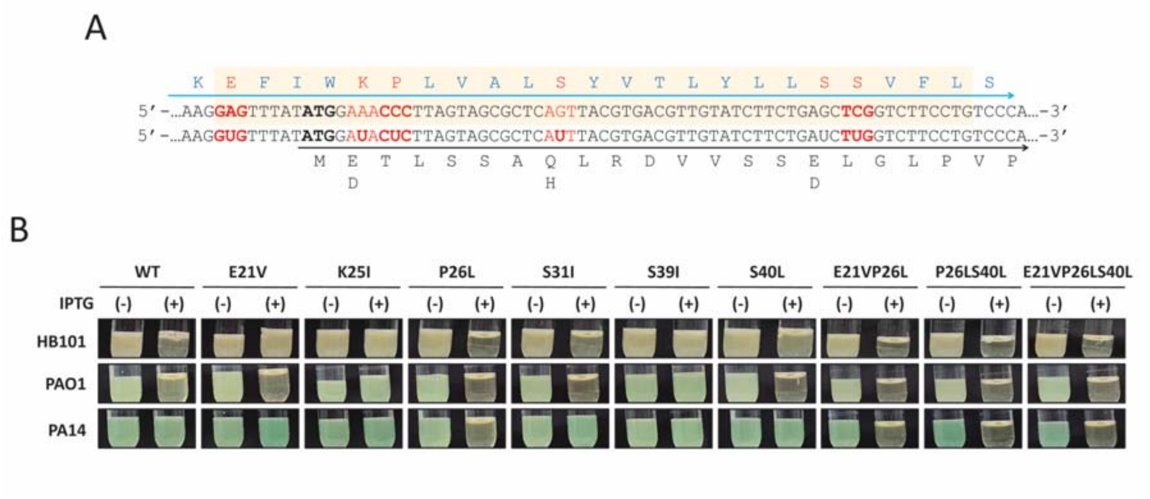


**Figure S4. Bacterial killing mediated by expression of TM helix mutants,** related to Figures 3 and 4

A. Mutation residues in LP. The partial sequences of the LP gene are shown with the translated LP (upper) and RNA replicase (RP) (lower). The codon changes are designated at the lower DNA sequence just beneath the WT DNA sequence. The codons for the 6 amino acids are shown in red with the mutations shown in bold. Three mutations (E21V, P26L, and S40L) do not change the amino acids in RP, whereas the other 3 mutations (K25I, S31I, and S39I) lead to the mutations in RP as designated (E2D, Q8H, and E16D, respectively).

B. Growth inhibition of the LP mutants. The six single mutants and the 3 P26L-derived mutants without codon changes in RP were grown in 15-ml tube format liquid culture for 6 h with (+) or without (-) IPTG induction. PAO1(LP) was included as the control designated as WT. The growth inhibition was monitored by visual inspection.

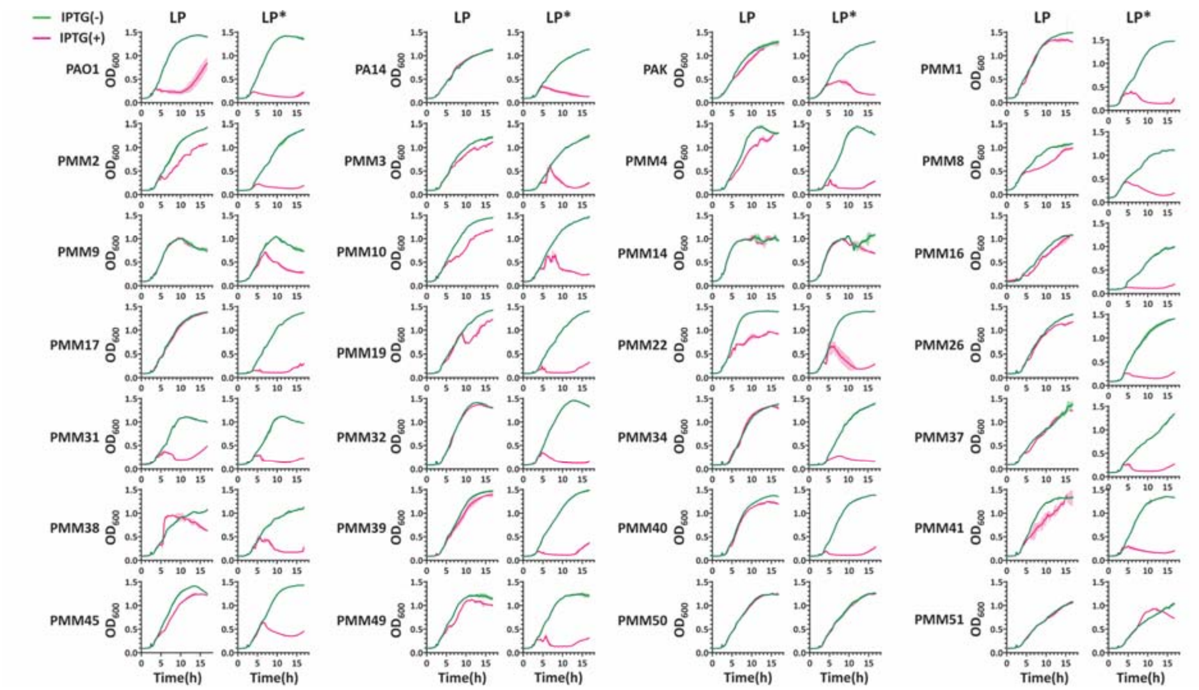


**Figure S5. LP-mediated growth inhibition of PA clinical strains,** related to Figure 5

Twenty-eight PA strains including PAO1, PA14, and PAK, with the expression system for either LP or LP* were grown in 96-well format liquid culture for 18 h with or without IPTG induction at 140 min and the growth was monitored by measuring the OD600 every 20 min for 17 h. PAO1(LP) and PAO1(LP*) cells harboring the MucP expression plasmid (pUCP-mucP) are included as the positive controls.
